# PPARG in osteocytes is essential for sclerostin expression, bone mass, marrow adiposity and TZD-induced bone loss

**DOI:** 10.1101/2020.09.13.295378

**Authors:** Sudipta Baroi, Piotr J. Czernik, Amit Chougule, Patrick R. Griffin, Beata Lecka-Czernik

## Abstract

PPARG role in regulation of osteocyte function is largely unknown. We report that PPARG is essential for sclerostin production, a recently approved target to treat osteoporosis. There is an excellent correlation in osteocytes between *Sost*/sclerostin and PPARG at the transcript and protein levels, and increased bone mass in mice with osteocyte-specific deletion of PPARG (γOT^KO^) correlated with increased WNT signaling and bone forming activity of endosteal osteoblasts and decreased marrow fat. The 8 kb sequence upstream of *Sost* gene transcription start site possesses multiple PPARG binding elements (PPREs) with at least two of them binding PPARG with dynamics reflecting its activation and the levels of *Sost* transcript and sclerostin protein expression. Older γOT^KO^ female mice are largely protected from TZD-induced bone loss providing proof of concept that PPARG in osteocytes can be pharmacologically targeted. Our study opens the possibility to consider repurposing PPARG as a target for treatment of osteoporosis.

## Introduction

Growing number of compelling evidences point to the mechanistic and functional connection between energy metabolism and bone mass (1–3). Peroxisome proliferator-activated receptor gamma (PPARG) exemplifies this connection. PPARG is a transcription factor that regulates network of genes associated with glucose and lipid metabolism, insulin signaling, adipocyte differentiation, and inflammation. PPARG has been implicated in the pathology of numerous diseases like diabetes, obesity, cancer, and atherosclerosis (4–6). In bone, PPARG controls osteoblasts and adipocytes differentiation from bone marrow stroma cells (BMSC) (7,8), and osteoclasts differentiation from cells of hematopoietic lineage, monocytes (9). Increased activity of PPARG in response to pharmacologic treatment with TZDs, PPARG full agonists and insulin sensitizers, results in bone loss associated with accumulation of fat in bone marrow, and up to two-fold increase in fracture rate in older women with diabetes (3,10,11). Activated PPARG increases osteoclast resorptive activities, and is skewing BMSC lineage allocation toward adipocytes and away from osteoblasts, which at least in part explains PPARG negative effects on bone. Our previous studies have indicated that there is an *in vivo* relationship between PPARG activity and *Sost* expression in osteocytes, and most importantly that the bone anabolic effect of a new class of insulin sensitizers, which act as PPARG inverse agonists and block S112 dephosphorylation, is associated with decreased *Sost* expression (12). These findings prompted us to explore the link between PPARG activity and sclerostin production with systematic and mechanistic approach.

Produced in osteocytes, the most abundant cells in bone, sclerostin protein regulates bone formation and resorption *via* inhibition of WNT signaling and production of RANKL (13). The newly approved anti-osteoporotic therapy with romosozumab consists of a monoclonal antibody targeting sclerostin. Romosozumab is highly efficacious with respect to the anabolic effect on bone and fracture prevention in postmenopausal women; however, there are safety concerns caused by the growing number of reported serious cardiovascular (CV) effects among the users of this drug, especially among individuals with high risk for CV events (14–18). Although the exact etiology of romosozumab-triggered CV effects is not determined, the off-target effects of the drug may include sclerostin produced in vascular endothelium and heart valves (19), and animal studies suggest that sclerostin produced by endothelium have a protective effect against vascular calcification (20). The growing list of potential, although not yet clinically recognized off-target effects, includes also joint diseases. It has been shown that sclerostin improves posttraumatic osteoarthritis (21) and that sclerostin produced by synovial fibroblasts have an inhibitory effect on progression of the TNFα-dependent rheumatoid arthritis, which is exacerbated with anti-sclerostin antibody therapy (22). Based on the above concerns and the superior anti-osteoporotic effects of anti-sclerostin therapy, the transcriptional control of sclerostin production in osteocytes can be a great alternative approach to the antibody therapy, especially when the transcription factor has been already identified as a therapeutic target, which is a case for PPARG.

Thus, with our long-term objective to demonstrate the feasibility of therapeutic targeting of PPARG to treat simultaneously diabetes, diabetic bone disease, and osteoporosis we designed current study to understand the role of PPARG in regulation of osteocyte function. To achieve this goal, we have developed mouse model of osteocyte-specific deletion of PPARG using *Dmp1*Cre driver and *Pparγ* loxP system. Here, we report that PPARG in osteocytes is a viable target to regulate sclerostin levels specifically in bone, and as a consequence bone mass and marrow adiposity, and skeletal response to pharmacologic activators of PPARG. Sclerostin is transcriptionally regulated by PPARG and PPARG is essential for sclerostin expression. These findings support consideration of selective pharmacological targeting of PPARG in osteocytes to control sclerostin production.

## Results

### *Pparγ* is highly expressed in osteocytes, and its expression increases with aging and with rosiglitazone treatment

*Pparγ* expression was analyzed in femora cortical bone fractions highly enriched in either osteocytes or endosteal osteoblasts, as validated at the level of cell type specific gene markers expression (Suppl. Fig.S1). As shown in Fig. 1A, an expression of *Pparγ* in osteocytes isolated from 6 mo old C57BL/6 males was 8-fold higher than the expression in endosteal osteoblasts isolated from the same femora cortical bone, whereas in 6 mo old females osteocyte expression of *Pparγ* was 40-fold higher than in osteoblasts. Interestingly, *Pparγ* mRNA expression increases with aging, as osteocytes derived from 10 mo old females have 3-fold higher *Pparγ* levels than osteocytes derived from 6 mo old females (Fig. 1B). Finally, *Pparγ* mRNA expression increases in osteocytes of mice treated with TZD rosiglitazone (Fig. 1C) indicating that PPARG protein in osteocytes is functional, because it responds to activation with the high affinity ligand by autoregulating its own expression, a phenomenon well documented in cells of adipocytic lineage (4).

**Figure 1.**
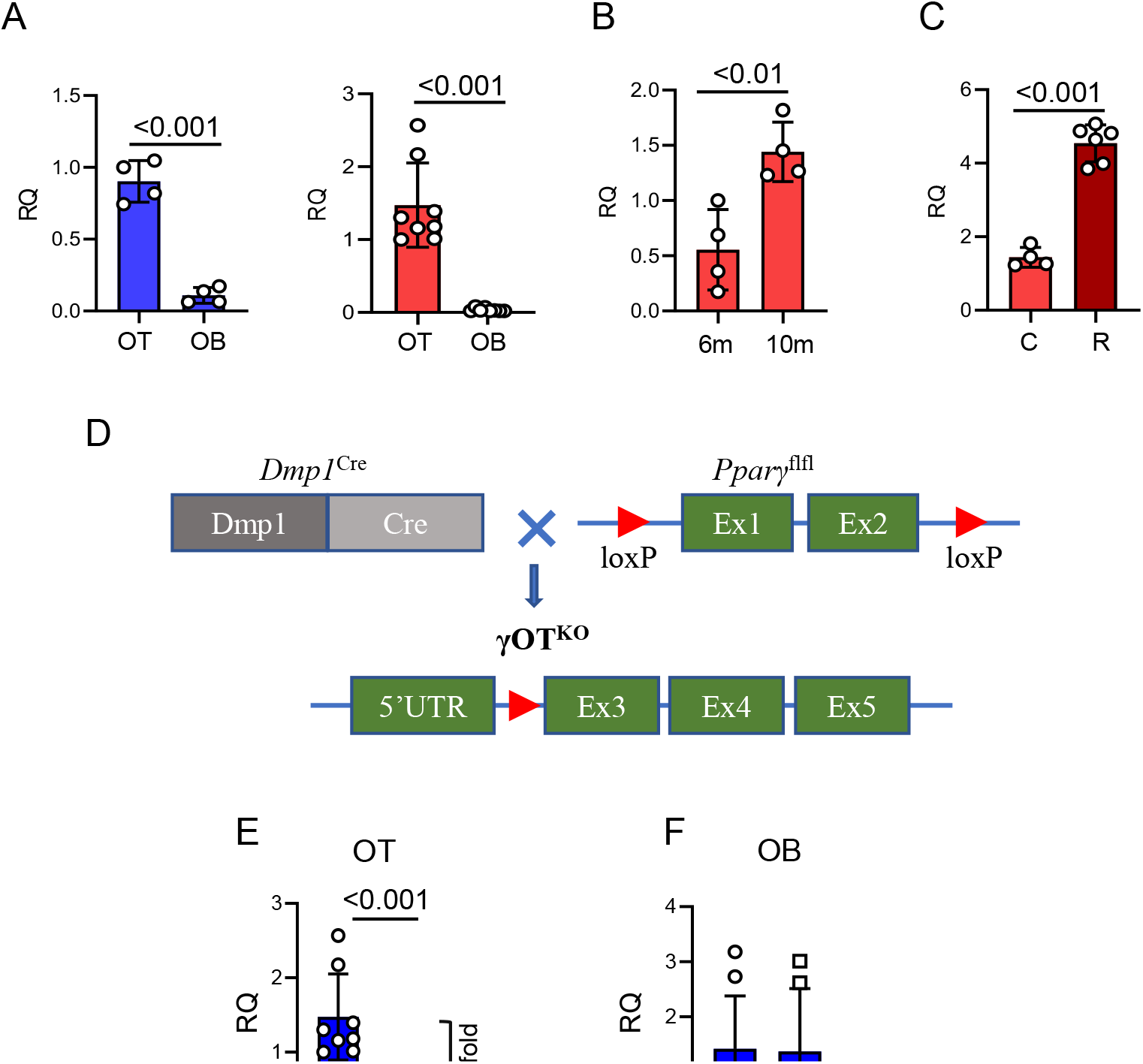
Development of γOT^KO^ mice. A. *Pparγ* is highly expressed osteocytes (OT) as compared to osteoblasts (OB). OT and OB were isolated from femora cortical bone of 6.5 − 7 mo old C57BL/6 males (blue) (γOT^KO^ n=4 and Ctrl n=4) and females (red) (γOT^KO^ n=8 and Ctrl n=8) using a method of differential collagen digestion and immediately processed for RNA isolation. B. *Pparγ* expression in OT increases with aging. OT were isolated as above from femora of 6 mo and 10 mo old C57BL/6 female mice. C. Rosiglitazone treatment increases expression of osteocytic *Pparγ*. OT were isolated from 10 mo old C57BL/6 female mice treated with 25 mg/kg/d rosiglitazone (R) for 6 weeks (γOT^KO^ n=6 and Ctrl n=4). D. Schematic of γOT^KO^ mice development. γOT^KO^ mice have deleted exon 1 and 2 from *Pparγ* gene, as a result of crossing Dmp1^Cre^ and *Pparγ*^flfl^ mice. E. *Pparγ* expression in OT isolated from 6 mo old γOT^KO^ (n=6) and Ctrl (n=8) male mice. F. *Pparγ* expression in OB isolated from the same mice as in E. Numbers above the horizontal bars indicate p values calculated with parametric unpaired Student ttest. RQ – relative quantity.

These findings gave us strong bases for seeking the role of PPARG nuclear receptor in osteocytes. Mice with PPARG deficiency primarily in osteocytes, the γOT^KO^ mice, were constructed by crossing *Pparγ*^flfl^ with 10kb *Dmp1*^Cre^ strains to remove exon 1 and exon 2 from the *Pparγ* gene, as shown in Fig. 1D. Offspring with a desirable γOT^KO^ genotype are viable and born with predicted Mendelian distribution with respect to sex and genotype. In all experiments, control mice (Ctrl) consist of littermates with either *Dmp*^Cre^*Pparγ*^WT^ or *Dmp*^WT^*Pparγ*^fl/fl^ genotypes to assure for a random distribution of parental genetic background.

The γOT^KO^ model was validated on several levels. The expression of *Pparγ* in osteocytes isolated from femora of γOT^KO^ mice was 100-fold lower than the expression in osteocytes isolated from Ctrl mice (Fig. 1E), while *Pparγ* expression in osteoblasts isolated from γOT^KO^ mice was not affected, as compared to Ctrl mice (Fig. 1F). Since there is a concern that Cre recombinase under control of 10 kb *Dmp1* promoter can be activated in other than osteocyte cell types, we analyzed the level of *Pparγ* expression in different tissues of adult γOT^KO^ and Ctrl mice. As shown in the Suppl. Table S1, we did not detect significant differences in *Pparγ* expression in muscle, bone marrow, liver, WAT, BAT, kidney, cerebellum, and small intestine of Ctrl and γOT^KO^ suggesting that even if 10 kb *Dmp1*^Cre^ construct is “leaky”, the expression of *Pparγ* in other tissues of adult γOT^KO^ mice is not obviously affected, which validates our study on the role of PPARG in osteocytes of mature bone.

### PPARG transcription factor is essential for sclerostin production

The level of PPARG protein deletion in osteocytes is paralleled by the level of sclerostin protein produced in these cells (Fig. 2A and 2B). There is an excellent correlation (R^2^=0.98) between PPARG and sclerostin levels, with both proteins being significantly decreased in osteocytes isolated from bone of γOT^KO^ mice (Fig. 2C). An analysis of sera of the same γOT^KO^ mice, show tendency to decrease in sclerostin levels but did not parallel a dramatic decrease in sclerostin protein in bone (Fig. 2D). These suggest that sclerostin levels in circulation are not in direct correlation with its production in bone and point to other sources for sclerostin production. Indeed, a number of studies indicates other organs such kidney, aorta and testes as producing sclerostin (23).

**Figure 2.**
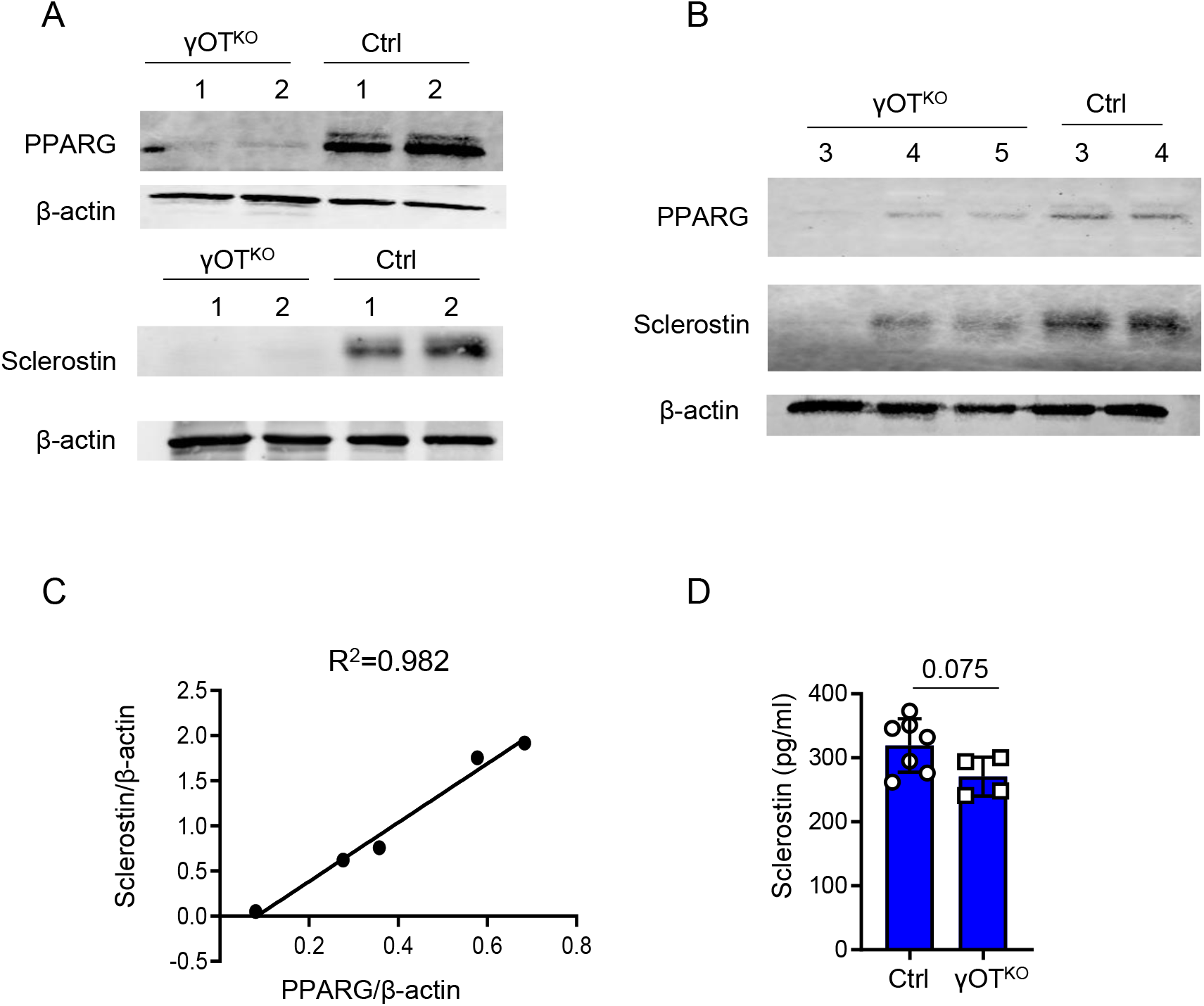
Correlation between PPARG and sclerostin levels in osteocytes. A. and B. Levels of PPARG KO and sclerostin proteins in OT isolated from 6.5 mo old male γOT^KO^ and Ctrl mice and either analyzed KO on separate (A) or the same (B) Western blot membrane. (γOT^KO^ n=5; Ctrl n=4). C) Pearson correlation analysis of PPARG and sclerostin protein levels shown in panel B. Coefficient of determination (R^2^=0.9821) was calculated based on bands density assessed by Image J and normalized to b-actin. D. KO Sclerostin levels measured by ELISA in sera of γOT^KO^ (n=4) and Ctrl (n=7) 6 mo old male mice. Statistical significance was calculated using parametric unpaired Student’s t test.

### Identification of PPREs located upstream of *Sost* transcription start site

The 8 kb DNA sequence upstream of transcription start site (TSS) of *Sost* gene was analyzed using a position weighted matrix (PWM) for prediction of binding sites for mouse PPARG::RXRA complex. The analysis was performed with assistance of the JASPAR database for curated, non-redundant set of transcription factor binding sites profiles. The position matrix algorithm is based on over 850 murine DNA sequences that were identified to bind the PPARG::RXRA heterodimer (Suppl. Fig. S2). Based on PWM, the JASPAR projected 15 PPARG response elements (PPRE) in the *Sost* gene promoter/enhancer region (Table 1 and Fig. 3A). The PPREs are located on both DNA strands. In the proximal 2 kb region, which represents *Sost* promoter, there are 4 PPREs, two of them with relatively low scores for PPARG binding consensus sequence are located within 1kb from TSS (PPRE1 and PPRE2) and the other two (PPRE3 and PPRE4) with higher scores at 1.8 – 1.9 kb distance from TSS. A cluster of 7 PPREs (PPRE5 to PPRE12) has been detected within 1 kb fragment at the 3.5 – 4.5 kb distance from TSS. Binding scores for PPREs in this cluster are high and varied from 8.1 to 12.5. The last cluster, with relatively high scores, consists of 3 PPREs (PPRE13 to PPRE15) and is located at the 7 – 8 kb upstream from TSS (Fig. 3A and Table 1). For further analysis, we selected two PPRE-containing regions based on their location and the highest scores for the consensus of PPARG binding sequence. The proximal PPRE3 located in the promoter region and two distal PPRE14 an PPRE15 located in the potential enhancer region. The consensus score for PPARG binding is 9.2 for PPRE3, and 15.3 and 9.6 for PPRE14 and PPRE15, respectively (Table 1).

**Table 1.**
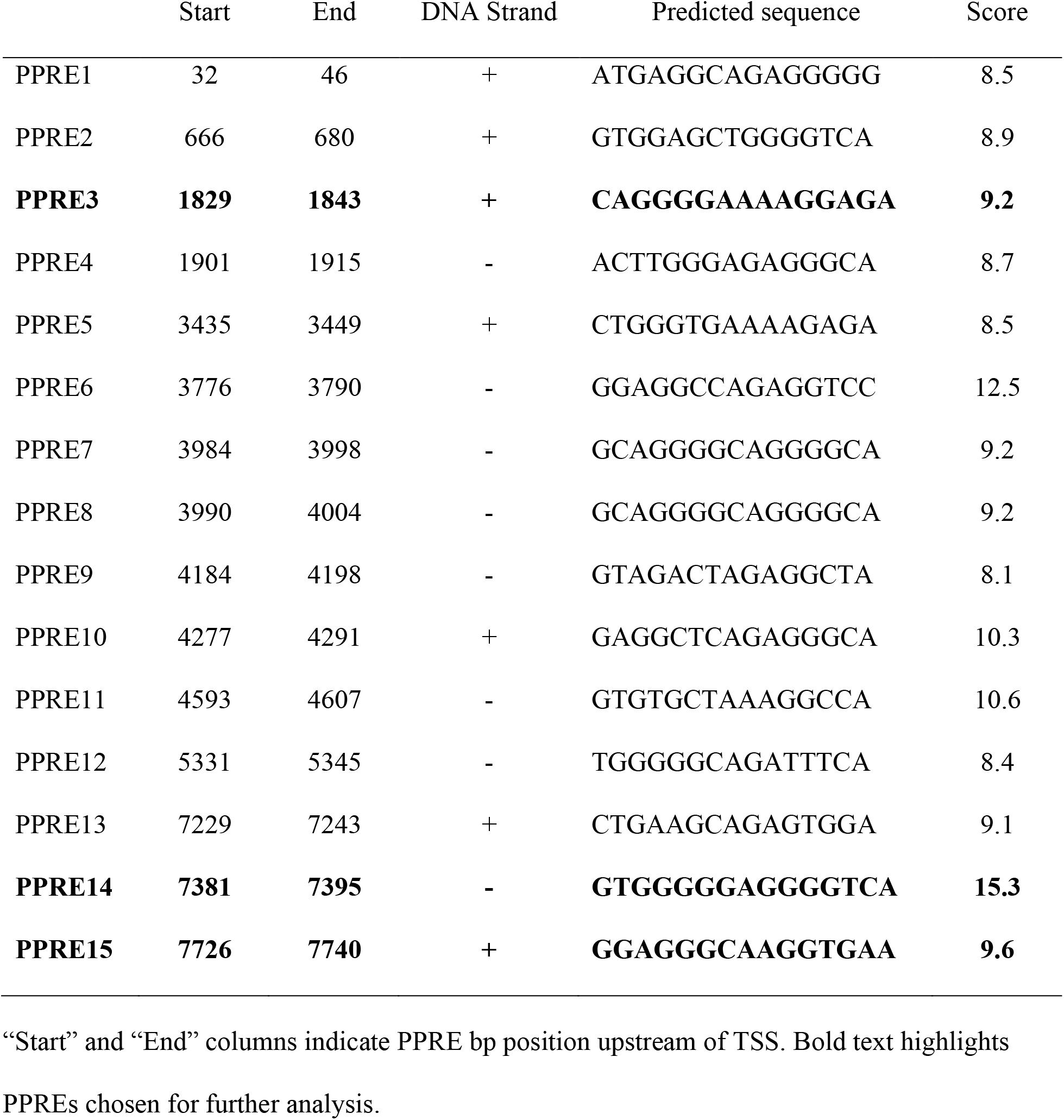
Predicted PPRE sequences identified within 8.85 kb sequence upstream to *Sost* gene TSS using JASPAR position weighted matrix

**Figure 3.**
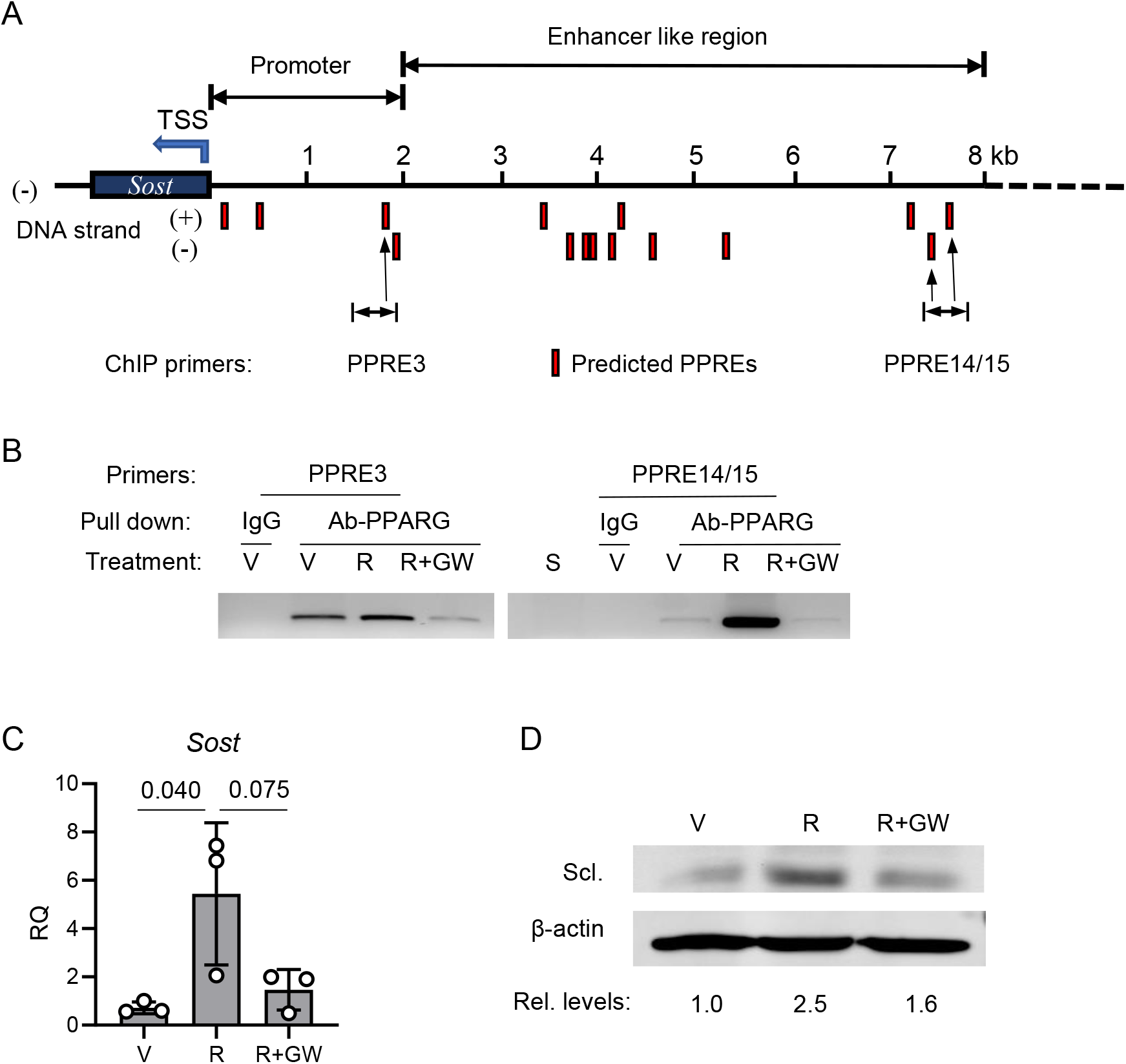
PPARG positively regulates sclerostin expression. A. Schematic positioning of JASPAR projected PPARG response elements (PPREs) upstream of sclerostin transcription start site (TSS). B. Chromatin immunoprecipitation (ChIP) assay end point agarose gel image (full image of the gel is presented in Suppl. Fig. 1). ChIP was performed on osteocyte-like MLO-Y4 cells targeting PPRE3 and PPRE14/15. Cells were treated for 24h with either vehicle (V), or 1µM rosiglitazone (R) or combination of 1µM R and 10µM GW9662 antagonist (R+GW) and ChIP assays were performed, as described in Material and Methods. S – sonicated lysate (no antibody pulldown). C. Expression of *Sost* mRNA in MLO-Y4 cells treated for 3 days with either vehicle (V), or rosiglitazone (R), or combination of rosiglitazone and GW9662 (R+GW) at the same doses as in (B). One-way ANOVA followed by Tukey’s post hoc analysis was performed for significance calculation. D. Western blot of sclerostin protein levels in MLO-Y4 cells treated as in (C). Sclerostin protein levels were normalized to β-actin levels measured as band density using Image J and relative levels of expression had been calculated and shown below Western blot images.

Using the chromatin immunoprecipitation assays (ChIP), we tested selected PPREs for their binding PPARG in basal conditions and upon activation with rosiglitazone. The assay was performed on osteocyte-like MLO-Y4 cells and showed that PPARG binds to both locations of the PPRE motifs (Fig. 3B). In basal conditions, PPARG protein occupies proximal PPRE3 and its binding to the PPRE14/15 location is rather weak. Activation with rosiglitazone slightly increases binding to PPRE3, while treatment with the covalent antagonist GW9662 significantly decreases PPARG binding to this site (Fig. 3B and Suppl. Fig. S3). In contrast, treatment with rosiglitazone induces robust recruitment of PPARG to PPRE14/15 binding sites, which is completely abrogated in the presence of GW9662 antagonist (Fig. 3B).

The pattern of dynamic changes of PPARG interactions with PPREs was paralleled by regulation of *Sost* mRNA expression in the same MLO-Y4 cells. Treatment with rosiglitazone upregulated *Sost* expression, which in the presence of GW9662 antagonist was attenuated (Fig. 3C). This was followed by changes in the sclerostin protein levels. Treatment with rosiglitazone increased sclerostin by 2.5-fold, which was then reduced to 1.6-fold in the presence of GW9662 antagonist (Fig. 3D). These results demonstrate that PPARG is a positive regulator of *Sost* transcription and sclerostin protein production. Most importantly, the PPARG transcriptional activity regulating sclerostin can be modulated pharmacologically.

### Expression of WNT signaling and bone formation markers is elevated in endosteal osteoblasts of γOT^KO^ mice, as compared to controls

Sclerostin acts as an inhibitor of WNT and BMP signaling in osteoblasts. Consistent with low levels of sclerostin in γOT^KO^ osteocytes, the expression of members of WNT signaling is upregulated in endosteal osteoblasts. Osteoblasts isolated from γOT^KO^ mice have increased expression of WNT ligands, *Wnt10b* and *Wnt16*, and β-catenin, the transcriptional mediator of canonical WNT pathway activity (Fig. 4A). This is followed by increased expression of genes, which are positively regulated by this signaling, including *Axin2*, *Connexin43* and *Cyclin D* (Fig. 4B). We have also analyzed expression of osteoblast-specific BMPs, *Bmp2* and *Bmp4*, and found that *Bmp4* expression is significantly increased in endosteal osteoblasts derived from γOT^KO^ mice (Fig. 4C). Finally, an increase in activity of WNT and BMP pathways is reflected in upregulation of osteoblast-specific transcription factors (*Runx2* and *Dlx5*) and alkaline phosphatase (*Alp*), a regulator of bone matrix mineralization (Fig. 4D).

**Figure 4.**
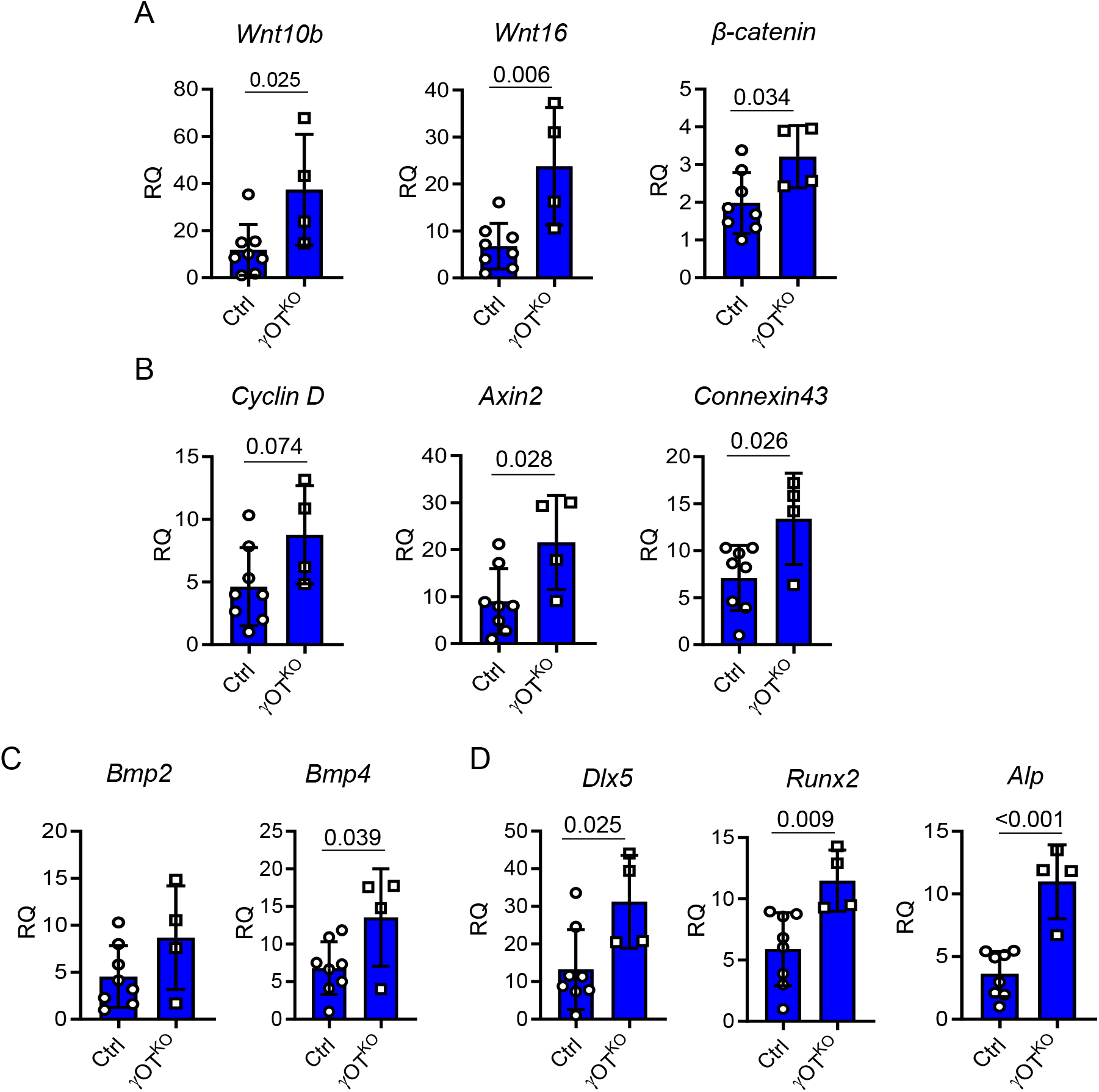
Relative expression of signaling pathways and osteoblast gene markers analyzed in fraction of endosteal osteoblasts freshly isolated by differential collagen digestion of femora cortical bone of 6.5 mo old males. A. WNT pathway signaling gene markers. B. Expression of genes positively regulated by WNT signaling. C. BMP signaling gene markers. D. Expression of genes positively regulated by WNT KO and BMP signaling and representing markers of bone forming osteoblasts. Ctrl (n=8) and γOT^KO^ (n=4). Statistical significance was calculated using parametric unpaired Student’s t test.

### γOT^KO^ mice have higher BMD and increased trabecular, but not cortical, bone mass

In support of increased WNT signaling and osteoblast activity, γOT^KO^ mice, both males and females, have high bone mass. In males, an increased global bone mineral density (BMD) is noticed as early as 4 mo of age, while in females it develops later in life (Fig. 5A). An analysis of bone formation rate in calcein double-labeled trabecular bone of proximal tibia showed that both γOT^KO^ males and females at the age of 6 mo have higher bone formation rate (BFR) as compared to Ctrl mice with intact PPARG in osteocytes (Fig. 5B).

**Figure 5.**
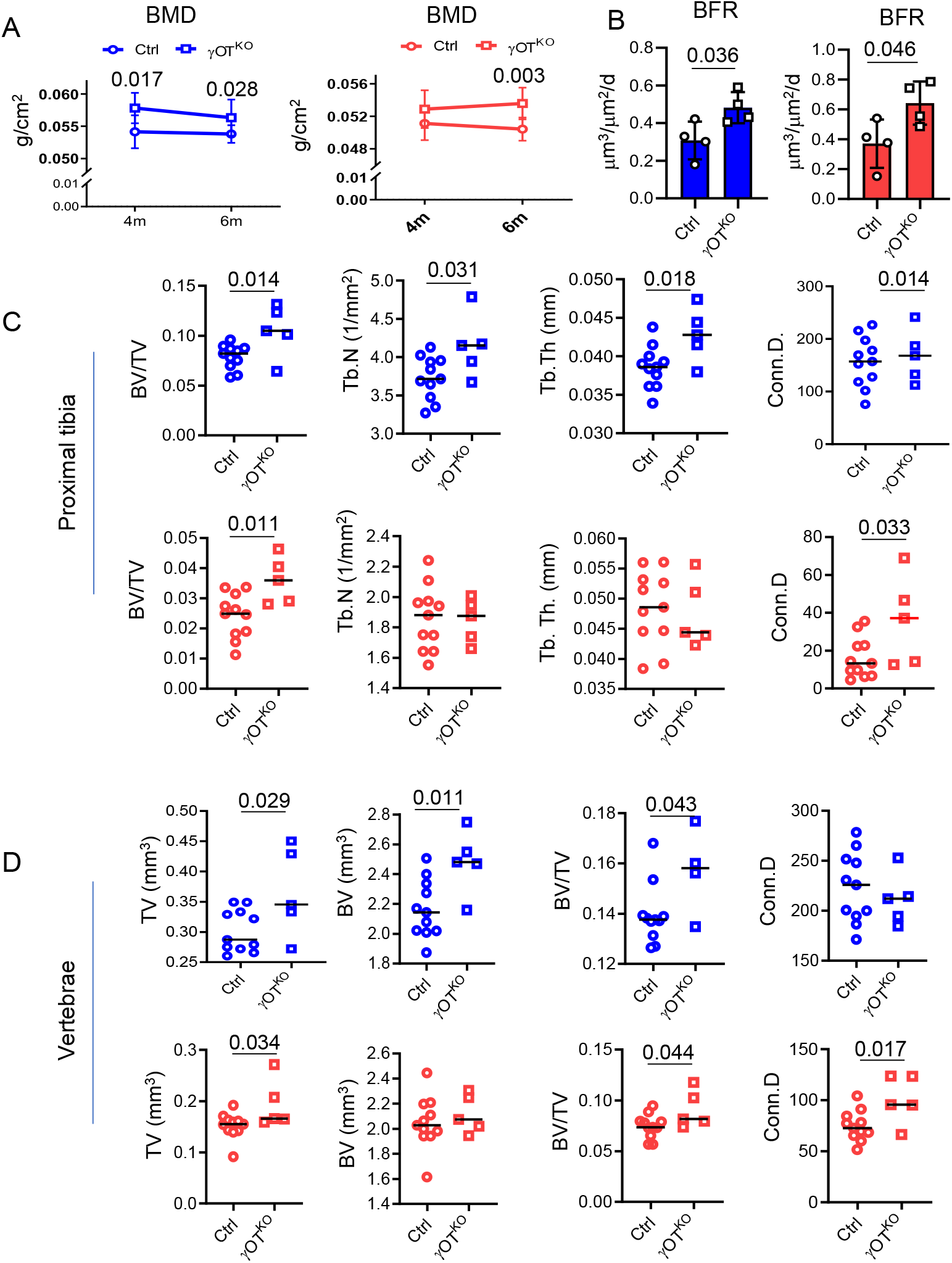
Both males (blue) and females (red) γOT^KO^ mice exhibit high bone mass and high bone formation. A. Measurements of global BMD using DXA at 4 mo and 6 mo of age. Ctrl (n=11) and γOT^KO^ (n=5). B. Bone formation rate (BFR) in 6.5 mo old males and 7 mo old females measured in calcein double-labeled tibia. Ctrl (n=4) and γOT^KO^ (n=4). C. µCT measurements of trabecular bone in proximal tibia. D. µCT measurements of trabecular bone in L4 vertebrae. C and D. Ctrl (n=11) and γOT^KO^ (n=5). TV – tissue volume, BV – bone volume, BV/TV – trabecular bone mass, Tb.N – trabecular number, Tb.Th – trabecular thickness, Conn.D – connectivity density. Statistical significance was calculated using parametric unpaired Student’s t test.

An analyses of bone microarchitecture using micro-computed tomography (mCT), showed that both sexes have increased trabecular bone mass in appendicular and axial skeleton (Fig. 5C and 5D). In males at 6 mo of age, high trabecular bone mass in proximal tibia was consistent with high trabecular number and thickness, whereas in females at the same age high trabecular bone mass correlated rather with structural changes at the level of connectivity density (Fig. 5C). Similarly, both males and females have high trabecular bone mass in L4 vertebra which correlated with increased connectivity density (Fig. 5D). Interestingly, in both sexes the vertebra body is larger in γOT^KO^, as compared to Ctrl mice (Fig. 5D). Remarkably, the changes in global BMD and trabecular bone mass were not followed by changes in properties of tibia cortical bone. The cortical thickness, bone area, marrow area, and bone material properties did not differ in γOT^KO^ and Ctrl regardless of sex (Suppl. Table S2). The skeletal phenotype of γOT^KO^ mice is consistent with a phenotype of mice with sclerostin ablated under 10kb *Dmp1* Cre control, which are characterized with high trabecular bone mass and an absence of effect on cortical bone (24).

### PPARG in osteocytes regulates marrow fat volume at least in part *via* sclerostin axes

Besides regulation of WNT signaling in osteoblasts, sclerostin is also implicated in regulation of marrow adipocyte differentiation, probably through the inhibitory effect on WNT pathway activity in cells of adipocyte lineage (25). In general, marrow fat content in tibia of γOT^KO^ mice is decreased. In males, the decrease in total volume of bone marrow adipose tissue (BMAT) is mostly due to its decrease in distal part since the proximal tibia contains very little fat in 6 mo old animals (Fig. 6A). However, in females, which have much more BMAT in proximal tibia than males, PPARG ablation in osteocytes resulted in substantial reduction of BMAT in this location (Fig. 6B and 6C). Similarly to males, females have decreased BMAT volume in distal tibia. Pearson correlational analysis of BMAT volume and osteocyte levels of sclerostin protein in the same animals resulted in the partial but statistically significant coefficient of determination, with the value R^2^ = 0.695 (Fig. 6D).

**Figure 6.**
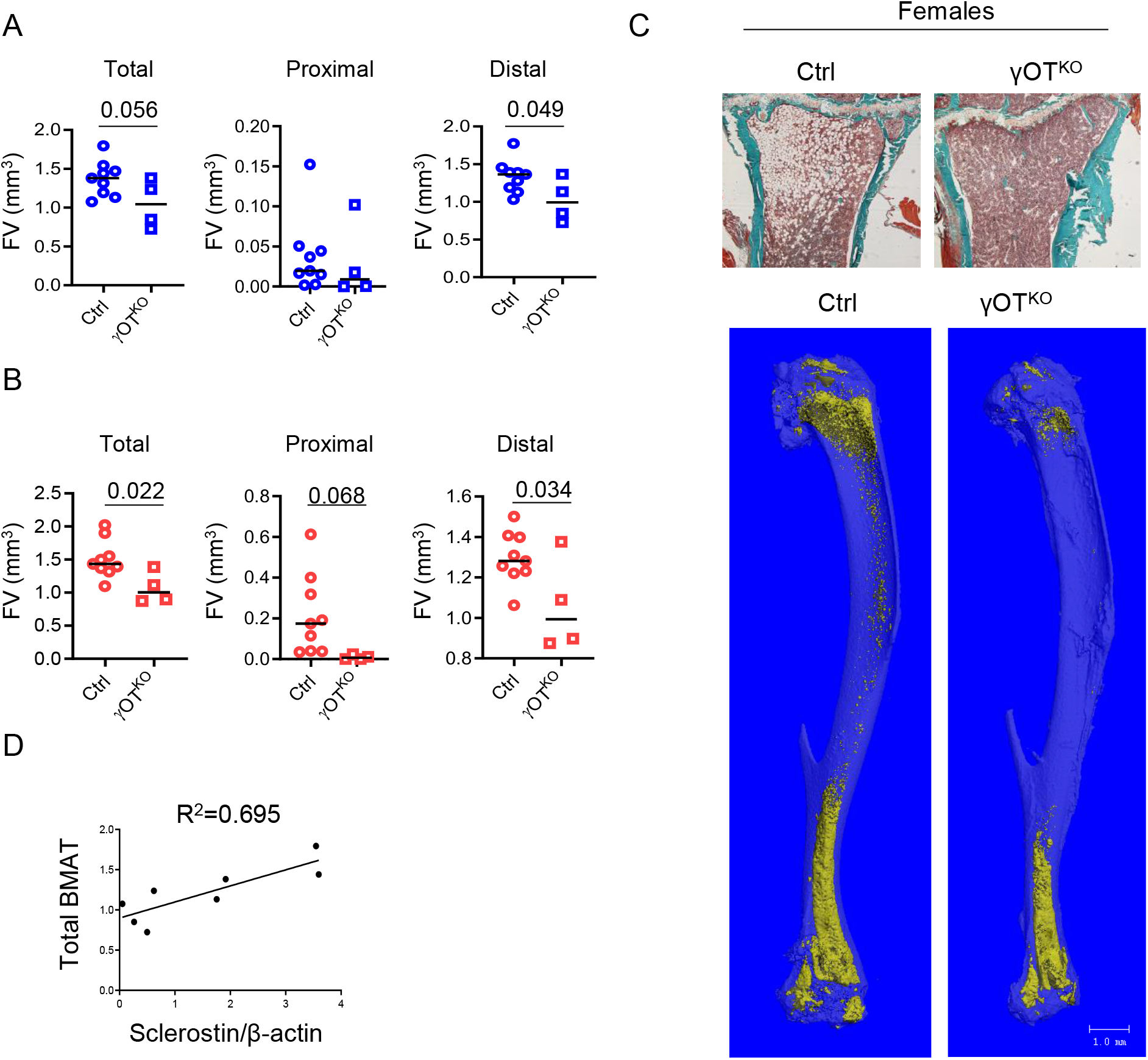
PPARG in osteocytes regulates marrow fat content. A. BMAT content in tibia of 6.5 mo old males γOT^KO^ (n=4) and Ctrl (n=9) mice. Measurements were done using µCT after decalcification of tibia followed by OsO_4_ staining. B. BMAT measurements in tibia of 7 mo old females γOT^KO^ (n=4) and Ctrl (n=9) mice. C. Representative Goldner’s Trichrome stained proximal tibia of female Ctrl and γOT^KO^ mice and renderings of OsO_4_ stained entire tibia (below) µCT images of female Ctrl and γOT^KO^ mice. Bar on mCT renderings indicates 1 mm. D. Pearson correlation of marrow fat volume and PPARG protein levels in osteocytes of Ctrl and γOT^KO^ mice (males, n=8). Statistical significance was calculated using parametric unpaired Student’s t test.

The coculture experiment assessed sclerostin contribution to the regulation of BMSC lineage allocation toward adipocytes (Fig. 7A). Conditioned media, collected from primary cultures of osteocytes isolated from femora of intact C57BL/6 mice and depleted from sclerostin with specific antibodies (Fig. 7B), decreased expression of *Adiponectin* and *Fabp4*, and have a tendency to decrease an expression of adipocyte-specific *Pparγ2,* in recipient BMSC isolated from the same mice (Fig. 7C).

**Figure 7.**
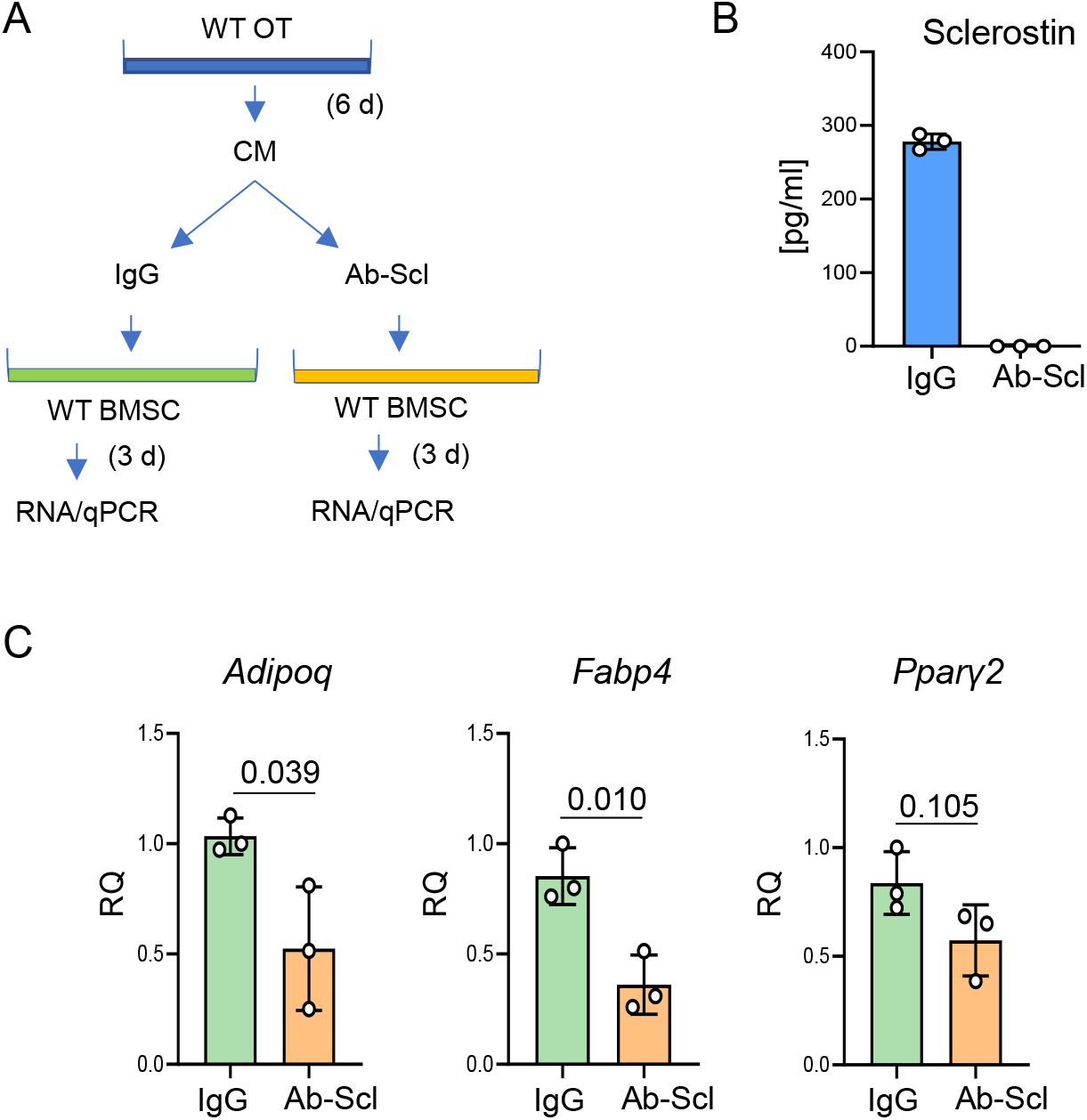
Osteocyte derived sclerostin positively contributes to marrow adipocyte differentiation. A. Schematic showing experimental design of co-culture of BMSC with intact or sclerostin depleted conditioned medium (CM). One group of adherent BMSC culture received IgG depleted CM from primary osteocytes (control group) while the other group received sclerostin depleted CM (group of interest). B. ELISA measurements of sclerostin level in CM after anti-SOST antibody mediated depletion. C. Expression of adipocytic and osteoblastic gene markers in adherent BMSCs treated with CM from primary osteocytes IgG or sclerostin (Ab-Scl) depleted. Statistical significance was calculated using parametric unpaired Student’s t test.

A significant decrease in BMAT volume in γOT^KO^ mice, together with a partial correlation between sclerostin levels and BMAT volume, and a decrease in adipocytic phenotype of BMSC subjected to CM depleted from sclerostin, indicate that PPARG in osteocytes is essential for physiological BMAT formation and that sclerostin contributes to this process.

### γOT^KO^ mice are partially resistant to rosiglitazone-induced bone loss

The evidence presented in Fig. 1C and Fig. 3, collectively imply that PPARG activated with rosiglitazone increases sclerostin production. It has been documented extensively by our lab and others that rosiglitazone decreases bone mass in mice and humans. Therefore, we asked a question of a contribution of osteocyte PPARG to the skeletal effects of rosiglitazone. Female mice at the age of 10 mo were fed a diet supplemented with rosiglitazone for 4 weeks. As shown in Fig. 8A, a global bone mineral density (BMD) decreased substantially in Ctrl mice as a result of treatment, while BMD of γOT^KO^ mice did not change. However, mCT examination of trabecular bone mass in vertebra body showed a tendency to decrease in γOT^KO^ mice. While BV/TV of Ctrl mice decreased significantly by 34.4%, the vertebra BV/TV of γOT^KO^ mice decreased by 20.7% although did not achieve statistical significance (Fig. 8B). More detailed examination of trabecular microarchitecture confirmed that PPARG in osteocytes contributes to a great extend to the bone loss due to rosiglitazone treatment. While in Ctrl mice rosiglitazone significantly decreased trabeculae number and increased spacing between trabeculae, the γOT^KO^ mice are largely protected from these effects, although some tendency to changes in these parameters maybe observed. An analysis of trabecular bone in distal femur was unsuccessful due to the very low trabeculae bone fraction (approx. 1%) left in this location in Ctrl older female mice and we did not detect significant changes in femora cortical bone which had been previously reported in male mice (26).

**Figure 8.**
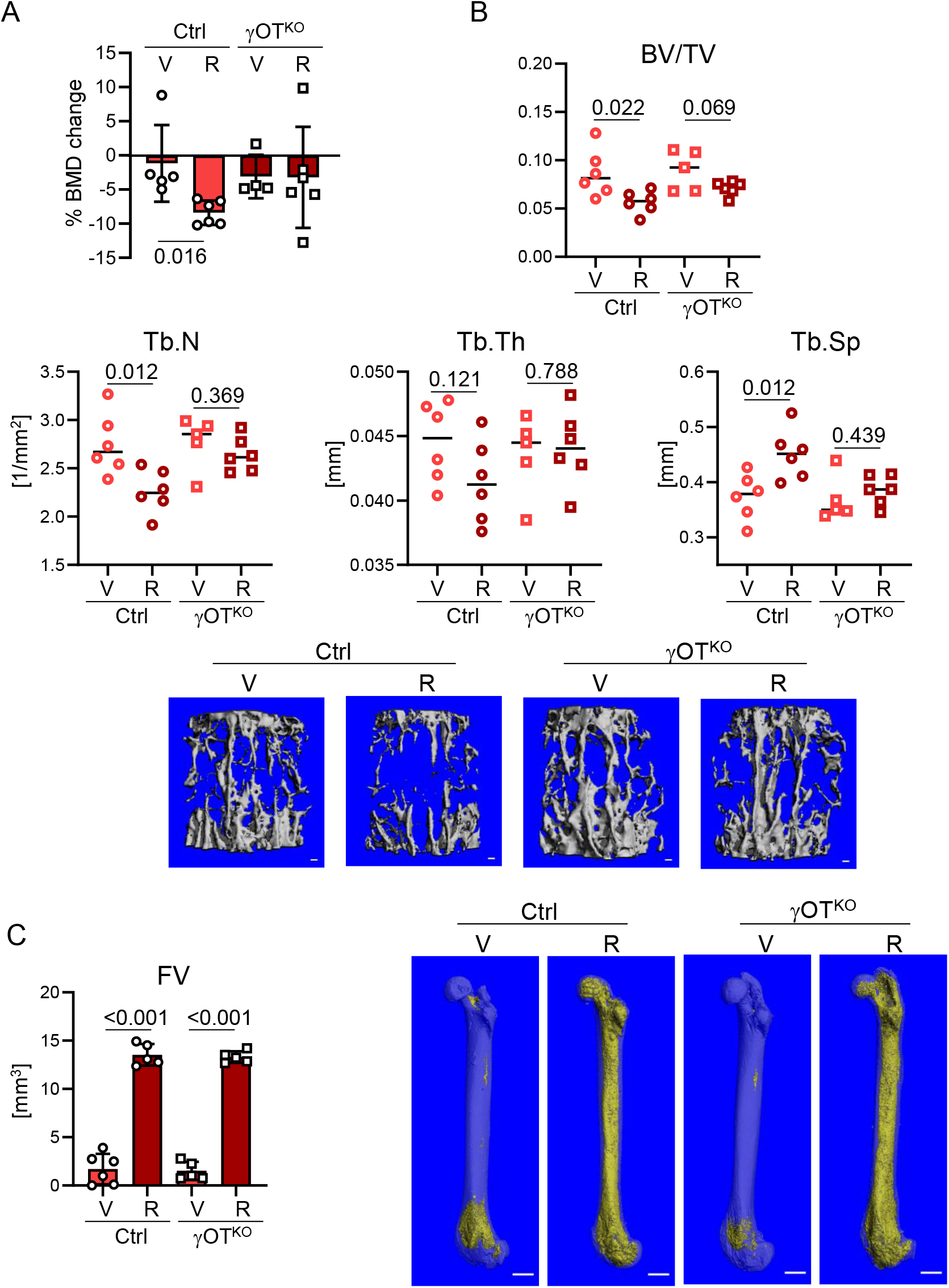
Bone of γOT^KO^ mice is partially protected from the detrimental effects of rosiglitazone. A. Percent change in global BMD after 4 wks treatment of 10 mo old females with rosiglitazone (25 mg/kg/d). n=4-6/group. B. µCT measurements of trabecular bone in L4 vertebrae and representative renderings. n=5-6/group. BV/TV – trabecular bone mass, Tb.N – trabecular number, Tb.Th – trabecular thickness, Tb.Sp – trabecular spacing. Bar on renderings indicates 0.1 mm. C. Left – Total fat volume in femora measured with µCT after decalcification and staining with OsO_4_ (n=5/group). Right – Representative mCT renderings. Bar on renderings indicates 1 mm. Significant p values are indicated over horizontal lines, statistical significance of rosiglitazone effect was calculated using parametric unpaired Student’s t test.

Surprisingly, PPARG deficiency in osteocytes did not protect from accumulation of large quantities of marrow fat in response to rosiglitazone treatment (Fig. 8C). These results indicate that TZDs-related bone loss rely on PPARG function in osteocytes and may include sclerostin regulation, while proadipocytic effect of TZDs is separate and probably consists of direct effect on BMSC, which in γOT^KO^ model have unaffected expression of PPARG protein. This also means that PPARG in osteocytes does not play a significant role in rosiglitazone-induced BMAT accumulation.

## Discussion

Presented studies provide compelling evidence that PPARG in osteocytes plays an important role in transcriptional regulation of sclerostin expression. First, we have shown that there is an excellent positive correlation between protein levels of PPARG and sclerostin in osteocytes. It is notable that in the absence of PPARG there is an absence of sclerostin, which is pointing to this nuclear receptor as essential for sclerostin production. Consequently, PPARG activation with full agonist rosiglitazone increases *Sost* transcript expression and sclerostin protein levels. Second, we have identified multiple PPRE sequences upstream of transcription start site, and demonstrated that at least two of them bind PPARG protein with dynamics reflecting status of PPARG activation. Thus, PPARG on molecular levels actively interacts with *Sost* gene DNA regulatory elements located in promoter/enhancer fragment upstream of TSS. Third, an ablation of PPARG in osteocytes leads to increased bone mass and decreased physiological levels of BMAT. Increases in the global BMD and trabecular bone mass, which are associated with increased WNT signaling in endosteal osteoblasts, are expected outcome of sclerostin deficiency in osteocytes (13,24). Finally, PPARG deficiency in osteocytes mitigates rosiglitazone-induced bone loss to a large degree, but does not prevent massive accumulation of fat in marrow cavity, indicating that osteocytes are the major contributors to the TZD-induced bone loss, but not to the medically-induced marrow adiposity. Although, we cannot conclude with certainty that the skeletal resistance to rosiglitazone treatment is entirely mediated by a lack of sclerostin, however presented results provide proof of concept that pharmacologic modulators of PPARG activity reach osteocytes in their lacunae and exert the function including differential regulation of sclerostin protein.

Finding the functional link between PPARG and sclerostin is of great importance, because both proteins are pharmacologic targets for existing therapies to treat diabetes and osteoporosis, respectively. This finding provides means to explore therapeutic overlap between anti-diabetic and anti-osteoporotic therapies. In fact, our previous study showed that anabolic effect on bone of a novel type of insulin sensitizers, which act as inverse agonists in respect to the PPARG pro-adipocytic and anti-osteoblastic activity, was associated with down-regulation of *Sost* expression in osteocytes (12). Here, we have demonstrated that PPARG is a positive regulator of sclerostin protein expression by showing that its ablation from osteocytes shuts down, while its activation increases, sclerostin production.

A potential prospect to pharmacologic targeting PPARG to reduce sclerostin in bone have several caveats which need to be corroborated. Does PPARG regulate sclerostin in other organs too, including aorta, kidney or joints? These are potential off-targets for therapies with sclerostin antibodies (14–18,20,22). Interestingly, we have shown that deletion of PPARG followed by an absence of sclerostin in osteocytes does not dramatically change the levels of sclerostin in circulation. This corresponds to the observation of others that there is a weak correlation between the levels of sclerostin production in osteocytes and the levels in circulation, indicating a substantial contribution of other organs to the total sclerostin levels (23,24,27). Thus, paracrine effect of sclerostin produced in osteocytes directly affects endosteal osteoblasts, with a little, if any, contribution of circulating sclerostin. Our results are supporting this notion. The lack of sclerostin production in osteocytes results in increased bone mass and activation of endosteal osteoblasts, even in the presence of relatively unchanged sclerostin levels in circulation. However, it needs to be addressed whether regulation of sclerostin expression in other organs is also under the control of PPARG. An existence of multiple PPRE sequences in the *Sost* promoter region may indicate tissue-specific regulation, and may hold a premise for differential modulation of PPARG activities controlling sclerostin production specifically in osteocyte.

Since discovery of sclerostin and its encoding gene *Sost*, there have been great efforts to identify bone-specific transcriptional mechanisms regulating its expression. The transcriptional control of *Sost* is complex and implicates number of signaling pathways and only few identified transcription factors, which directly bind to the promoter region and regulate its activity. Several signaling pathways have been recognized as important for *Sost* expression, the activities of which converging on either the promoter region or enhancer elements of *Sost* gene. Among them, the most prominent is PTH signaling, which acts on ECR5 enhancer *via* MEF2c/HDAC5 transcriptional regulators, and negatively regulates *Sost* expression in osteocytes (28–30). However, there is ample of evidence for other signaling mechanisms acting either in a positive (BMP, TNFα, Vitamin D, IL-1α, IL-1β, TWEAK, RA, and calcitonin), or a negative (PGE2, Oncostatin M, Cardiotrophin, and Sirtuin1), or both manners determined by biological context (TGFβ, glucocorticoids, estrogen, and hypoxia), and interacting with either an immediate promoter region or more distal enhancers (31,32)).

Mechanistically, activities of these signaling are channeled to the interaction of specific transcriptional regulators with cognate regulatory elements in the *Sost* promoter/enhancer regions. Up to date, few of them have been identified for their binding to the promoter including RUNX2, OSX/SP7, ZFP467, and TIEG (33–36). An array of loss- and gain-of-function experiments determined that RUNX2, OSX/SP7 and ZFP467 act as positive regulators of *Sost* promoter activity and that at least between RUNX2 and OSX/SP7, which binding sites are located in the proximal promoter region between −106 bp to −260 bp upstream of TSS, there is a great degree of coordination in this regulation (37). In contrast, TIEG, or TGFβ inducible early gene-1, has been identified as a negative regulator of sclerostin expression acting downstream of estrogen signaling (36). Interestingly, two identified KLF/SP1 binding sites for TIEG, which are located at −1864 kb and −1956 kb upstream of TSS, are in close proximity to the PPRE3 (−1829 kb), identified here as binding PPARG, and PPRE4 (−1901 kb). It is well recognized that PPARG and estrogen signaling oppose each other during adipose tissue expansion, including BMAT expansion in conditions of estrogen deficiency (38–40). It also has been shown that estrogen is a negative regulator of sclerostin (36). Thus, there is a prospect that PPARG interaction with distal region of *Sost* promoter may be affected by, or may be subjected to, the modulatory effect of neighboring regulatory elements, such as KLF/SP1 perhaps in estrogen dependent manner. On the other hand, two distal PPRE14 and PPRE15, which have very high scores for PPRE consensus sequence and bind PPARG with dymamics coresponding to the status of PPARG activition, are located approximately in the 1 kb distance upstream to the region identified as enhancer responsive to Vitamin D and possessing a putative VDR binding element (41), which creates a possibility of PPREs interaction with a modulatory element under control of another nuclear receptor, VDR.

On note, we have also identified up to 7 PPREs with relatively good scores, which are clustered within a 1 kb fragment at the 4 kb distance from TSS; however, in this study we did not analyze the functional significance of this region to the regulation of *Sost* expression. Nevertheless, the pattern of such close clustering is known to facilitating strong interaction with binding proteins, even if the particular consensus sequences are not at the highest scores (42). We have previously identified similar pattern of PPREs distribution in the promoter of genes known to be directly and positively regulated by PPARG including adipocyte-specific gene *Fabp4* (Supplementary Fig. 4 in ref. (8)). The significance of this region to *Sost* gene expression remains to be established for both PPARG basal activity and the activity modulated pharmacologically by either full or inverse agonists. Taking into account the PPRE number, pattern of their distribution, and binding activity of those which were analyzed, it concludes in a strong support for PPARG directly regulating *Sost* expression *via* interaction with promoter and enhancer region.

Besides regulating bone remodeling, osteocytes also modulate marrow environment including marrow adiposity with sclerostin having a prominent role in this process. By the fact that sclerostin acts as an inhibitor of WNT pathway activity, it has a role in skewing BMSC lineage allocation toward adipocyte and away from osteoblast differentiation. The proadipocytic activity of sclerostin has been reported in several systems of adipocyte differentiation including 3T3-L1 cell line representing progenitors for white adipose tissue (43), ear mesenchymal stem cells, and BMSC (44). Consistent with proadipocytic activity, a decrease in sclerostin protein levels in γOT^KO^ mice correlates with a decrease in BMAT volume. However, there are few interesting aspects of this phenotype including different manifestation in respect to sex and skeletal location, and a degree of correlation.

Several research groups including our have shown that BMAT represents heterogenous tissue consisting of adipocytes originating from different progenitors and having different phenotype and function depending on skeletal location and exposure to environmental or hormonal signaling (40,45,46). Here we show that ablation of PPARG in osteocytes affects BMAT in tibia to a different degree in respect to location and animal sex. We observe an approximately 20% decrease in the volume of distal BMAT regardless of animal sex. However, proximal BMAT in females is affected to much larger degree in γOT^KO^ mice indicating of powerful signals derived from osteocytes that regulate BMAT formation. The coculture experiments with intact or sclerostin depleted conditioned media confirm that sclerostin is partially, but not entirely, responsible for this phenotype. The degree of inhibition of adipocyte gene markers in recipient cells *in vitro* together with a rather modest correlation of BMAT volume in tibia with sclerostin levels in the same bone, suggest that there are other factors under PPARG control in osteocytes, which regulate BMAT specifically in the proximal tibia location in females. We did not observe proximal BMAT phenotype in male tibia, probably because at the age of 6 mo they did not have sufficient amount of BMAT in this location that would allow for non-biased analysis. Nevertheless, the differences in BMAT volume in proximal and distal tibia indicate that proximal BMAT is to a large degree dependent on osteocyte signaling under control of PPARG.

The significance of presented study is reinforced by the fact that there is a scarcity of information on the role of PPARγ in osteocytes. To our knowledge, there are only two reports that previously addressed this issue. The first consists of development a model of PPARG deficiency in osteocytes constructed from the same *Dmp1*Cre driver and *Pparγ*flfl mice, which was characterized with high bone mass including increased cortical bone mass and periosteal bone formation; the phenotype that we did not confirm in our model. This study was limited to male mice and did not focus on the mechanism by which PPARG in osteocytes controls bone mass including regulation of sclerostin expression and bone marrow composition in respect to BMAT (47). The second model, demonstrated that sclerostin expression is increased in osteocyte-like MLO-Y4 in response to treatment with PPARG agonist rosiglitazone, which is consistent with our results; however, the mechanism of this effect was not provided (48).

In presented study, we have focused on the PPARG-sclerostin connection; however, we recognize that this relationship is a part of a larger role which PPARG plays in regulation of osteocyte function supporting bone homeostasis and energy balance; the role which is pursued in another ongoing study. The current study indicates that PPARG is an essential regulator of osteocyte function with substantial fraction of these activities being channeled through sclerostin activities.

## Materials and Methods

### Animals

Osteocyte-specific PPARG knock-out mice, γOT^KO^, were developed at the University of Toledo according to the Institutional Animal Care and Utilization Committee protocol. The γOT^KO^ mice were constructed by crossing the 10kb *Dmp1*^Cre^ mice (B6N.FVB-Tg (*Dmp1*- cre)1Jqfe/BwdJ) and *Pparγ*^fl/fl^ mice (B6.129-*Pparg*^tm2Rev^/J). Both mouse strains have C57BL/6 background and were obtained from the Jackson Laboratory (Bar Harbor, ME). *Pparγ*^fl/fl^ mice have loxP sites flanking exons 1 and 2 of *Pparγ* gene. The Cre-recombinase is under late osteoblast/osteocyte specific10kb *Dmp1* promoter. The desired osteocyte-specific γOT^KO^ and littermate control mice were obtained in the F3 progeny of 10 kb *Dmp1*^Cre^ and *Pparγ*^fl/fl^ cross. As controls, mice with either *Dmp1*^Cre^ only and/or *Pparγ*^fl/fl^ only genotypes were used. Animals were housed under 12h dark-light cycle with unlimited access to chow (Teklad global 16% protein rodent diet; code 2916) and water. During breeding of these mice, breeding diet (Teklad global 19% protein extruded rodent diet; code 2919) was used.

For genotyping, tissue samples were collected at weaning using tail clipping and genomic DNA isolated after proteinase K (Sigma-Aldrich, Cat#P5568) digestion. PCR primers published by the Jackson Laboratory were used for genotyping. The Cre recombinase insertion was detected with following primers: F – TTG CCT TTC TCT CCA CAG GT and R – CAT GTC CAT CAG GTT CTT GC); while to differentiate between *Pparγ*^fl/fl^ (homozygous), *Pparγ*^fl/+^ (heterozygous) and *Pparγ*^+/+^ (wildtype) strains following primers were used: F – TGGCTTCCAGTGCATAAGTT and R – TGTAATGGAAGGGCAAAAGG. Taq polymerase (Go Taq; Promega, Cat#M3005) based amplification was used for PCR followed by 1% agarose gel run for amplicon detection (167 bp for Cre; 200 bp for *Pparγ* wildtype, 230 bp for *Pparγ* homozygotes, both 200 bp and 230 bp for *Pparγ* heterozygotes).

Rosiglitazone supplemented diet experiment: Nine month old female mice (Ctrl and γOT^KO^) were fed with either regular or rosiglitazone supplemented at the dose 25 mg/kg/day diet, as described (26). DXA was performed at the beginning of the study and after 4 weeks of treatment to measure changes in BMD. After 6 weeks of treatment, animals were sacrificed, and tissues harvested. Right femur and vertebrae were scanned with mCT for bone parameters, then femur was decalcified, stained with OsO_4_ and scanned for marrow fat.

### DXA

Dual Energy X-ray absorptiometry (DXA) was used to measure BMD values longitudinally as the mice age. Measurements were done using Lunar PIXImus II instrument (GE Lunar Corp., Madison, WI) at 4 mo and 6 mo of age and analyzed using LUNAR PIXImus v. 2.10 software provided by manufacturer. Head and metal tag region were excluded from region of interest (ROI).

### mCT analysis of bone and marrow fat

Right tibiae and L4 vertebrae were collected from euthanized mice and stored in 10% buffered formalin until scanned using µCT35 system (Scanco Medical AG, Basserdorf, Switzerland). A minimum 48h wait period was maintained before scanning to attain fixation plateau for all bones. mCT scans were performed at 70 peak kilovoltage (kVp) energy and 114- μA intensity settings and using 7-μm voxel. For proximal tibiae trabecular bone, integration time was 300 ms and 200 slices were analyzed, for midshaft cortical bone, integration time was 100 ms and 57 slices were analyzed, while for vertebrae, integration time was 100 ms and 100 slices were analyzed, as recommended (49).

To visualize marrow lipids, bones were decalcified using Formical 4 (Medline; Cat# STLB12145) and stained with freshly prepared 2% aqueous solution of Osmium Tetroxide (Electron Microscopy Sciences, Cat#19150) in 0.1 M sodium cacodylate buffer (pH 7.4), as described (50). Staining was carried out in an exhaust hood and away from light due to osmium tetroxide toxicity and light sensitivity. Images of lipid depositions were acquired at 70-kVP and 114-μA settings and 12-μm nominal resolution. Image segmentation was done under global threshold conditions by applying a gray scale threshold of 480–1000 using the per mille scale with the three-dimensional noise filter set to sigma 1.2 and support 2.0. Lipid volumes were calculated directly from individual voxel volumes in three-dimensional reconstructions.

### Dynamic histomorphometry

Sterile filtered Calcein (Sigma, Cat#C0875-5G) solution (2.5mg/ml) prepared in isotonic 2% NaHCO3 was administered intraperitonially (20mg/kg body weight) to mice, 10 days and 2 days before euthanasia. Left tibia was collected in 70% EtOH and embedded in methyl methacrylate (Indiana University Histology Core), and processed for slide preparation and staining. From the slides, images were taken using Nikon Eclipse 80i, for the proximal tibia region adjacent to growth plate. From the images, single and double calcein labelled surfaces were measured using Nikon NIS-Elements BR3.0 in a blinded manner by single operator. Endo-osteal bone surface in proximal tibia excluding growth plate surface was considered for measurement. Mineralizing surface (MS/BS), Mineral apposition rate (MAR), Bone formation rate (BFR) were calculated, as recommended (51).

### Organ culture conditioned medium experiment

Conditioned media were collected from organ cultures of femora bone. Femora from six C57BL6 mice were collected, bone marrow was spun out and discarded, the outer surface of bone was cleaned off any remaining muscle fiber and periosteum, and one collagenase digestion followed by one EDTA digestion was performed, as described in (52). Using sterile bone-cutting scissors, the bones were then chopped into 1-2 mm fragments and distributed evenly on collagen coated 6-well plates with 3 ml culture medium/well (αMEM supplemented with 15% FBS and 10 mM glucose). Bone pieces were cultured for 6 days at 37C with 5% CO_2_ before conditioned medium (CM) was collected and pooled from all six bone organ cultures. Sclerostin was depleted from one half of the collected osteocyte CM by overnight incubation with anti-sclerostin antibody (Anti mouse sclerostin Ab, polyclonal Affinity purified; R & D Systems; Cat# AF1589) in the ratio 20 µl antibody per 9 ml CM, and pulled down with Protein A agarose beads (Invitrogen, Cat#15918014). The other half of the conditioned media was subjected to IgG pull-down step (2 µl for 9 ml CM) (Anti-mouse IgG monoclonal Ab; Sigma-Aldrich; Cat#M9144) which was used as a negative control for this assay. Sclerostin levels after depletion were quantified using SOST ELISA kit (Quantikine ELISA, R&D Systems, Cat#MSST00).

Recipient BMSC cultures were established from bone marrow isolated from femora of six C57BL6 mice, pooled and plated on 6-well plated at the density 2×10^6^ cells/well. After 7 days of growth, the culture medium was then replaced with osteocyte CM either depleted of Sclerostin or IgG in triplicates, and cultured for three more days before processing for RNA isolation followed by analysis of adipocytic and osteoblastic markers using qPCR.

### Isolation of osteocyte-enriched and endosteal osteoblast-enriched cell fractions from femora

An osteocyte-enriched fraction and endosteal osteoblast enriched fraction after sequential collagenase digestion of femora (52,53). Briefly, femora were cleaned off muscles and bone marrow was spin-out after removal of epiphysis. The diaphysis-outer-surface was scrapped to remove periosteum and any remaining muscle fibers. Bones were then subjected to alternating digestion with collagenase and EDTA to retrieve endosteal osteoblasts and osteocytes. First two collagenase and EDTA digestions of femora were discarded. Cells from the 3^th^ digestion were collected and represented an endosteal osteoblast enriched fraction 3 (F3). Alternating digestions continued and cells from the 6^th^ digestion were collected and represented osteocyte enriched fraction 6 (F6). The leftover bone pieces were homogenized and added to F6 osteocyte enriched fraction. Cell-type enrichment of isolated fractions was verified by analyzing levels of osteoblasts (*Col1*, *Ocn*) and osteocytes (*Dmp1*, *Sost*) gene markers expression (Suppl. Fig. S1).

### Gene Expression Analysis

mRNA was isolated using Trizol (Thermo Fisher Scientific, Cat# 15596026). Half μg RNA was converted to cDNA using the Verso cDNA synthesis kit (Thermo Fisher Scientific, Cat#AB1453B). Gene transcripts were detected by quantitative real-time PCR using TrueAmp SYBR Green qPCR SuperMix (Smart Bioscience, Cat#7501-3) and processed with StepOne Plus System (Applied Biosystems, Carlsbad, CA). Relative gene expression was measured by the comparative ΔΔCT method using *18S* RNA levels for normalization. Primers were designed using OligoPerfect Designer (Thermo Fisher Scientific) and are listed in the Suppl. Table S3.

### Protein analysis using Western blots

Proteins were isolated from cell fraction enriched in osteocytes using TRIzol, following manufacturer’s protocol and quantified using BCA assay (Pierce BCA protein Assay Kit, ThermoScientific, Cat#23225). Ten µg of protein was loaded on each lane and resolved on 10% SDS-PAGE using BioRad system and electrophoretically transferred to Immobilon-FL membranes. Membranes were blocked at room temperature for 2h in Odyssey blocking buffer (LI-COR, Cat#927-50000). Incubation with primary antibody was performed overnight at 4° with either mouse monoclonal anti-PPAR*γ*, 1:500 dilution, (Santa Cruz Biotechnology, Santa Cruz, CA; cat # Sc-7273), or goat polyclonal anti-sclerostin, 1:500 dilution, (R&D System, Minneapolis, MN; cat #AF1589), or mouse monoclonal β-actin, 1:10,000 dilution, (Sigma-Aldrich Life Science, St. Louis, MO; cat #A1978). After incubation, membranes were washed 3 times in TBST (TBS plus 0.1% Tween 20), and incubated with appropriate secondary antibody, either infrared anti-mouse (680RD; Cat#925-68072) or infrared anti-goat (800CW; Cat#925-32214) (LI-COR Bioscience, Lincoln, NE), at 1:10,000 dilution in TBS for 2h at room temperature. Immunoreactivity was visualized by infrared scanning in the Odyssey system (LI-COR Biosciences). Image J (version 1.52a) was used to calculate band intensity from Western blots and band intensity of beta-actin was used to normalize PPARG and sclerostin signals.

### In-silico mapping of PPREs in Sost regulatory region using JASPAR

The 8.85 kb sequence upstream to *Sost* gene transcription start site (TSS) was obtained from NCBI Genebank database. This sequence was divided into 3 equal fragments (2.95 kb each) and analyzed for mouse PPARG::RXRA heterodimer binding sites using JASPAR (version 2020), a database for curated, non-redundant set of transcription factor binding profiles. Position weight matrix (PWM) for mouse PPARG::RXRA binding site was run with relative profile score threshold set at 80%.

### ChIP assay

Chromatin Immunoprecipitation (ChIP) assay to detect PPARG binding to selected PPRE sequences was performed on osteocyte like MLO-Y4 cells (kind gift from Dr. L. Bonewald). Cells were cultured on collagen coated 100 mm plates using 5% FBS, 5% CS and 1% P/S in alpha-MEM medium (at 37C with 5% CO_2_). Optimal passage time was 4-5 days before each splitting with 1:5 ratio after cultures reached 80% confluency. In each experiment, cells were treated in duplicates with either DMSO as vehicle, PPARG agonist rosiglitazone (1 μM) and a combination of rosiglitazone (1 μM) and antagonist GW9662 (10 μM). Sonicated only and Immunoglobulin G (IgG) pull-down samples were used as negative controls.

Chromatin Immunoprecipitation protocol was followed (54). Briefly, after 24h treatment cells were fixed using 4% formaldehyde in PBS, lysed, and DNA was sonicated to fragments size 500 – 1000 bp (6 min total time per sample, 12 cycles of 30s pulse at 50% amplitude of Qsonica CL-18 sonicator followed by 90s incubation on ice after each cycle). After sonication, samples were incubated with Protein A agarose beads (Invitrogen, Cat#15918014) for 4h at 4C to reduce background. The beads were spun down and discarded, and either IgG (2 μl/sample; Anti-mouse IgG monoclonal Ab; Sigma-Aldrich; Cat#M9144) or PPARG antibody (10 μl/sample; Anti-mouse PPAR*γ* monoclonal Ab; Santa Cruz Biotechnology;Cat#Sc-7273) were added and incubated overnight at 4C followed by adding fresh protein A agarose beads (60 μl/sample) and incubated for 4h at 4C. After spinning down the beads, antibody bound chromatin-PPARG fragments were eluted using elution buffer (54) followed by crosslink reversal (15h at 65C), Proteinase K digestion (1h at 50C) and purification of pulled-down DNA fragments using phenol-chloroform-isoamyl alcohol (25:24:1) extraction. The Taq polymerase (GoTaq) based PCR was performed with specifically designed primers for analyzed PPREs: PPRE3 F – TCTCCCAAGTCTGGAGCAAT and R – CTGAGGTGCAAAAGGAGGAG; PPRE14 & 15 F – TACCACGTCTCCCCTGTTTC and R – GGGCCTGTGTTTGCATAGTT. The presence of specific amplicons (532 bp for PPRE3 and 634 bp for PPRE14/15) was detected using 1% agarose gel.

### Statistical analysis

Data are represented as means ± SD and were analyzed by statistical analysis software GraphPad Prism 8. Parametric, unpaired, two-tailed Student’s t-test was used for comparing means of two groups. One-way ANOVA followed by Tuckey’s test (as post-hoc analysis) was used to compare multiple groups. Pearson correlation was used to measure correlation between experimental variables. A p-value 0.05 and lower was considered as statistically significant.

## Acknowledgements

This work was supported by grants from the National Institute of Diabetes, Digestive and Kidney Diseases (NIDDK) (R01 DK105825) to PRG and BLC, and the American Diabetes Association Innovative Basic Science Award (1-19-IBS-029) to BLC.

## Authors’ roles

SB and BLC contributed to experimental design, data analysis, and data interpretation; SB, PJC, and AC conducted the experiments; SB, BLC, and PRG wrote and edited the manuscript, SB and BLC take responsibility for the integrity of the data analysis.

