## Supplemental Material for "PPARG in osteocytes is essential for sclerostin expression, bone mass, marrow adiposity and TZD-induced bone loss"

Supplementary Table S1. Comparison of $C_T$ values of *Dmp1* and *Pparγ* mRNA expression in different tissues of γOT$^{KO}$ and Ctrl mice.

| Tissue | Genotype | *18S* | *Dmp1* | | | *Pparγ1* | | | *Pparγ2* | | |
|---|---|---|---|---|---|---|---|---|---|---|---|
| | | $C_T$ | $C_T$ | $\Delta C_T$ | $\Delta\Delta C_T$ | $C_T$ | $\Delta C_T$ | ΔΔCT | $C_T$ | $\Delta C_T$ | $\Delta\Delta C_T$ |
| Muscle | Ctrl | 11.68 | 36.72 | 25.19 | 2.37 | 32.69 | 21.01 | 1.45 | 32.94 | 21.26 | 0.98 |
| | γOT$^{KO}$ | 11.85 | 34.81 | 22.96 | 5.74 | 33.19 | 21.34 | 1.14 | 33.29 | 21.45 | 0.85 |
| Kidney | Ctrl | 10.14 | ND | | | 31.73 | 21.58 | 0.86 | 33.46 | 23.32 | 0.58 |
| | γOT$^{KO}$ | 10.60 | ND | | | 32.18 | 21.58 | 0.88 | 33.65 | 23.05 | 0.56 |
| Cerebellum | Ctrl | 10.59 | 28.77 | 18.17 | 0.64 | 31.07 | 20.48 | 1.05 | 32.49 | 21.90 | 0.75 |
| | γOT$^{KO}$ | 10.58 | 29.42 | 18.84 | 0.64 | 31.42 | 20.84 | 1.23 | 32.70 | 22.12 | 0.55 |
| Small intestine | Ctrl | 10.19 | 34.64 | 24.45 | 0.66 | 30.76 | 20.57 | 0.98 | 32.77 | 22.58 | 0.71 |
| | γOT$^{KO}$ | 10.12 | 34.95 | 24.83 | 0.64 | 30.27 | 20.15 | 1.23 | 32.88 | 22.76 | 0.55 |

$C_T$ values of real time PCR represent average of $C_T$ Mean from Ctrl (n=4) and γOT$^{KO}$ (n=3) male mice. No statistically significant differences were observed in the expression of *Dmp1*, *Pparγ1*, and *Pparγ2* in analyzed tissues between Ctrl and γOT$^{KO}$ mice. Muscle consists of gastrocnemius and soleus combined. ND – transcript not detected.

Supplementary Table S2. mCT cortical bone analysis

| | Males | | | | | Females | | | | |
|---|---|---|---|---|---|---|---|---|---|---|
| | γOTKO | | Ctrl | | | γOTKO | | Ctrl | | |
| | Mean | SD | Mean | SD | p value | Mean | SD | Mean | SD | p value |
| T.Ar ($mm^2$) | 2.033 | 0.259 | 2.066 | 0.131 | 0.730 | 1.558 | 0.120 | 1.617 | 0.102 | 0.327 |
| B.Ar ($mm^2$) | 0.722 | 0.116 | 0.691 | 0.031 | 0.411 | 0.553 | 0.046 | 0.570 | 0.031 | 0.415 |
| M.Ar ($mm^2$) | 1.311 | 0.154 | 1.375 | 0.119 | 0.374 | 1.005 | 0.076 | 1.048 | 0.087 | 0.363 |
| Ct.Th. (mm) | 0.194 | 0.011 | 0.190 | 0.008 | 0.460 | 0.182 | 0.011 | 0.185 | 0.009 | 0.523 |
| BMD (mg HA/$cm^3$) | 583.4 | 19.1 | 565.4 | 22.7 | 0.147 | 607.9 | 27.2 | 610.7 | 40.6 | 0.892 |
| TMD (mg HA/$cm^3$) | 1048.9 | 12.5 | 1047.8 | 7.8 | 0.828 | 1048.5 | 9.1 | 1055.2 | 25.4 | 0.583 |
| pMOI ($mm^4$) | 0.222 | 0.045 | 0.224 | 0.018 | 0.879 | 0.128 | 0.016 | 0.136 | 0.017 | 0.381 |
| Imax/Cmax ($mm^3$) | 0.180 | 0.027 | 0.176 | 0.011 | 0.670 | 0.111 | 0.014 | 0.115 | 0.012 | 0.623 |
| Imin/Cmin ($mm^3$) | 0.133 | 0.020 | 0.135 | 0.009 | 0.781 | 0.094 | 0.008 | 0.099 | 0.009 | 0.309 |

T.Ar – total area, B.Ar – bone area, M.Ar – marrow area, Ct.Th – cortical thickness, BMD – bone mineral density, TMD – tissue mineral density, pMOI – polar moment of inertia or torsional strength, Imax/Cmax - bending strength across the bone along the maximal centroid-to-edge distance, Imin/Cmin - bending strength across the bone along the minimal centroid-to-edge distance

Supplementary Table S3. Primers used for real time PCR

| Transcript | Forward | Reverse |
|---|---|---|
| *Pparγ1* | AGTGTGACGACAAGGTGAC | TGTTGGTCTCACAGGCTCC |
| *Pparγ2* | TGGGTGAAACTCTGGGAGATTC | TAGGCAGTGCATCAGCGAAG |
| *Dmp1* | TGTCATTCTCCTTGTGTTCCTTTG | AGAGCTTTCAGATTCAGTATTGTGGTAT |
| *Sost* | CCTCCTCCTGAGAACAACCA | ACATCTTTGGCGTCATAGGG |
| *Alp* | ATGGGCGTCTCCACAGTAAC | AGGGGAATTTGTCCATCTCC |
| *β-catenin* | TGCGGGAACAGGGTGCTA | TGCGCCGTTGGGTGTC |
| *Dlx5* | TGACAGGAGTGTTTGACAGAAGAGT | CGGGAACGGAGCTTGGA |
| *Runx2* | GGGCACAAGTTCTATCTGGAAAA | CGGTGTCACTGCGCTGAA |
| *Axin2* | TAG GCG GAA TGA AGA TGG AC | CTGGTCACCCAACAAGGAGT |
| *Wnt10b* | GTGCTTTCTCCTTCTCCATGC | CTCACCACTACCCTTCCATCC |
| *Wnt16* | GGAGCTGTGCAAGAGGAAAC | GAAGTGGTAGTGGCGACCAT |
| *Bmp2* | AACTGGCTAGAATATTAAGCACTGCA | AGTGATTTCCTAACTGCCCAGG |
| *Bmp4* | TCAAGGGAGTGGAGATTGGG | GCCATCATGGCCAAAAGTG |
| *Cyclin D* | CCAGAGGCGGATGAGAACAA | GGCACAGAGGGCCACAAA |
| *Adiponectin* | GGCCGTTCTCTTCACCTACG | TGGAGGAGCACAGAGCCAG |
| *Fabp4* | GCGTGGAATTCGATGAAATCA | CCCGCCATCTAGGGTTATGA |
| *Connexin43* | CAGGTGGACTGCTTCCTCTC | GAGCGAGAGACACCAAGGAC |

Supplementary Figure S1. Comparison of osteoblast ( *Col1A1* and *Ocn*) and osteocyte (*Dmp1* and *Sost*) gene markers expression in fraction 3 representing osteoblasts (OB) and fraction 6 representing osteocytes (OT) isolated with collagen digestion of cortical femora bone.

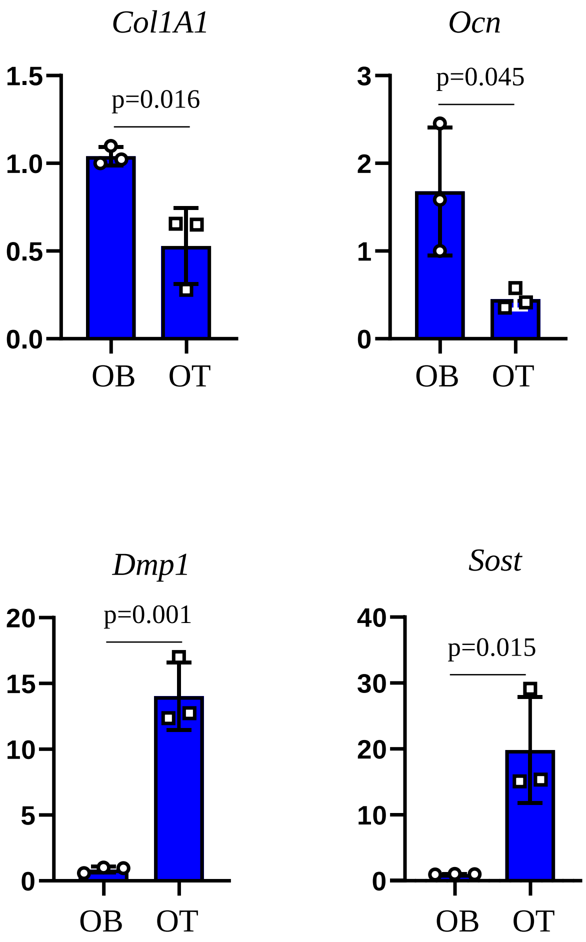

Supplementary Figure S2. Predicted position weighted matrix (PWM) of mouse PPARG::RXRA heterodimer response element according to Jaspar motif MA0065.2.

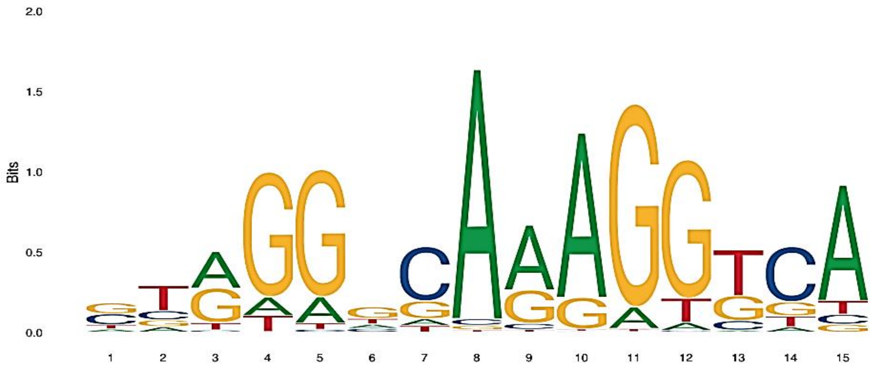


| Nucleotide position | A | C | G | T |
|---|---|---|---|---|
| 1 | 94 | 317 | 320 | 126 |
| 2 | 101 | 166 | 127 | 464 |
| 3 | 390 | 23 | 368 | 79 |
| 4 | 100 | 3 | 671 | 87 |
| 5 | 139 | 15 | 674 | 34 |
| 6 | 145 | 129 | 395 | 193 |
| 7 | 71 | 547 | 179 | 66 |
| 8 | 819 | 21 | 15 | 8 |
| 9 | 522 | 48 | 270 | 23 |
| 10 | 713 | 5 | 137 | 9 |
| 11 | 82 | 2 | 767 | 12 |
| 12 | 41 | 9 | 693 | 120 |
| 13 | 22 | 99 | 263 | 479 |
| 14 | 54 | 555 | 144 | 109 |
| 15 | 676 | 58 | 47 | 81 |

Supplementary Figure S3. Full image of agarose gel with PCR products from the representative ChIP assay.

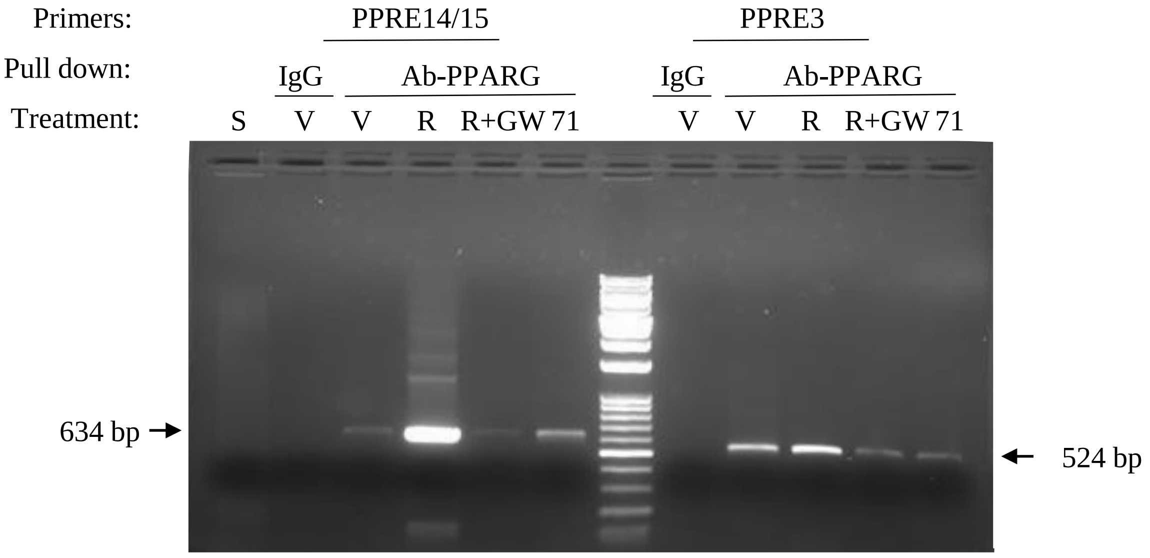


S-sonicated only; V-vehicle; R-Rosiglitazone; R+GW-Rosiglitazone + GW9662; 71 – SR10171

Band at 634 bp is specific for PPRE14/15 and band 524 bp for PPRE3
